# Caste, sex, and parasitism influence brain plasticity in a social wasp

**DOI:** 10.1101/2021.11.01.466692

**Authors:** Kristine M. Gandia, Federico Cappa, David Baracchi, Mark. E. Hauber, Laura Beani, Floria M. K. Uy

## Abstract

Phenotypic plasticity is the capacity of a single genotype to exhibit different phenotypes, and can be an adaptive response to specific environmental and social conditions. Social insects are particularly well-suited to study plasticity, because the division of labor amongst females and the different life histories of males and females are associated with specific sensory needs. Here, we take advantage of the social wasp *Polistes dominula* to explore if brain plasticity is influenced by caste and sex, and the exploitation by the parasite *Xenos vesparum*. Within sexes, males had proportionally larger optic lobes, while females, regardless of caste, had larger antennal lobes, which is consistent with sensory needs of sex-specific life histories. Within castes, reproductive females had larger calyces, as predicted by their sensory needs for extensive within-colony interactions and forming winter aggregations, than workers who spend more time foraging for nest material and prey. Surprisingly, parasites had different effects on female and male hosts. Female workers were castrated and behaviorally manipulated by female or male parasites, but only showed moderate differences in relative allocation of different brain tissue compared to non-parasitized workers. In contrast, the testes and behavior of parasitized males were essentially unaffected, but they had smaller brains and greater relative volume of most sensory brain regions than non-parasitized males. Our results are consistent with caste and sex mediating brain plasticity in *P. dominula* and that the parasite’s manipulation can also drive differential allocation of brain regions depending on host sex.

## INTRODUCTION

Within taxa, phenotypic plasticity generates responses to environmental variation, allowing for organisms with a single genotype to display a variety of adaptive phenotypes (West-Eberhard 2003; Whitman and Agrawal 2009). When plasticity optimizes energy allocation, it can result in tradeoffs between reproduction, growth, and allometric changes in body structures (Barton and Capellini 2011; Birget et al. 2017; Jokela and Mutikainen 1995). Brain plasticity, for instance, has evolved across many lineages as a result of variable selective pressures acting on the cognitive demands of sensory and perceptual systems (Barton et al. 1995; Barton and Harvey 2000; Catania 2005). Since brain tissue is energetically expensive, plasticity in specific brain structures may be linked to the sensory and processing needs of adaptive behaviors (Montgomery et al. 2016; Niven and Laughlin 2008; Riveros and Gronenberg 2010; Rozanski et al. 2021). Investment in neural tissue may be mediated by experience (Jones et al. 2009; Riveros and Gronenberg 2010), diet (Murphy et al. 2014), environmental stimuli (Burns et al. 2009), tradeoffs with reproduction (Pitnick et al. 2006), and/or endocrine factors (Lendvai et al. 2013). Therefore, understanding neuroanatomical plasticity can be challenging, as multiple factors may drive differential allocation of brain tissue (Willemet 2013).

Social insects, which have distinct caste systems with different life histories and sensory and reproductive needs (Beshers and Fewell 2001), provide the opportunity to compare plasticity in brain regions while controlling for genetic background. In insect brains, visual input is received and processed by the optic lobes, while olfactory input is received by the antennal lobes (Anton and Homberg 1999; Gronenberg and Hölldobler 1999; Strausfeld 1989). From these sensory neuropils, projection neurons convey the computed information to the mushroom bodies (Akalal et al. 2006). In these higher brain centers the chemical and visual information is further processed and integrated with internal information by intrinsic neurons and finally projected to premotor areas. In particular, the calyces of the mushroom bodies, acts as learning and memory centers and integrate this olfactory information in the lip, the visual information in the collar, and both sensory stimuli in the basal ring (Akalal et al. 2006; Ehmer and Hoy 2000; Fahrbach 2006). Finally, the central complex, is implicated in spatial navigation (Honkanen et al. 2019; Le Moël et al. 2019; Pfeiffer and Homberg 2014). Given that specialized behaviors in social insects are associated with a range of caste-specific sensory needs, corresponding investment in neural tissue is expected (Arganda et al. 2020; Ehmer et al. 2001; Gronenberg et al. 1996; O’Donnell et al. 2007; Penick et al. 2021; Rehan et al. 2015; Seid et al. 2011).

The primitively eusocial paper wasp *Polistes dominula* provides an excellent opportunity to test how brain plasticity is associated with behavioral flexibility (O’Donnell and Bulova 2017; O’Donnell et al. 2018; O’Donnell et al. 2014; Pardi 1996; Rozanski et al. 2021). In this temperate wasp species, the recognition of nestmates, caste, and sex relies on both chemical and visual cues (Beani et al. 2019; Cappa et al. 2016; Cappa et al. 2020; Cini et al. 2019; Dani et al. 2001). Females are morphologically similar and organized in a flexible caste system, according to a dominance hierarchy (Pardi 1948). The queen monopolizes egg-laying (Strassmann et al. 2004), while subordinate foundresses and workers are involved in nest building and defense, the rearing of larvae, and foraging (West-Eberhard 1969). The reproductive caste consists of gynes (future foundresses) and males which emerge in mid-summer. Gynes remain on the natal nest without performing any colony tasks and then mate, form winter aggregations, and enter diapause until the following spring (Reeve 1991). In contrast, males abandon the nest early after emergence, display lek-behavior at landmarks, mate with gynes (Cappa et al. 2013), and die at the end of summer (Beani 1996; Beani et al. 2014).

In addition, the intimate host-parasite relationship between *P. dominula* and the strepsipteran parasite insect *Xenos vesparum* provides a great opportunity to explore the effect of this parasite in allocation of brain tissue (Hughes and Libersat 2018; Libersat et al. 2018). *X. vesparum* manipulates the neuroendocrine physiology and behavior of females (Beani 2006; Hughes et al. 2004b; Strambi and Strambi 1973; Strambi et al. 1982). This parasite decreases the size of the corpora allata and castrates the females, by irreversibly inhibiting ovary development (Strambi and Strambi 1973). Parasitized workers abandon the colony and aggregate on selected plants where parasite mating occurs (Beani et al. 2018; Hughes et al. 2004b). Parasitized males, a secondary host, instead maintain their reproductive apparatus and sexual behavior (Beani et al. 2011; Cappa et al. 2014).

While brain plasticity within and across social insects has been extensively studied (Godfrey and Gronenberg 2019), to our knowledge, no studies have explored plasticity within a species that has morphologically similar individuals, various colony tasks, and a parasite that potentially alters brain morphology. We hypothesize that the relative volume of selected brain regions reflects specific sensory needs for each subset of wasps (reproductive females, female workers, and males) (Rozanski et al. 2021), and that parasitic manipulation may influence brain allometry. We predict higher volume of visual regions in males to detect and identify potential mates or rival males in a lek, compared to females. On the contrary, we expect more olfactory processing by females compared to males due to social interactions in the colony. We also predict a higher effect in brain plasticity towards parasitized workers, which are castrated and show aberrant behaviors, compared to parasitized males who reproduce and show no changes in behavior. Finally, we expand on studies that reported reduced corpora allata in parasitized females and males compared to non-infected conspecific by testing for the effect of parasite sex (Beani et al. 2017; Strambi and Strambi 1973).

## METHODS

### Field collection

We collected reproductive females (N = 10 foundresses and N = 9 gynes), along with non-parasitized workers (N = 10), workers parasitized by one *Xenos* female (N = 11) or by one *Xenos* male (N = 11) and males parasitized by one or two *Xenos* males (N = 9) during summers of 2016 and 2018, in the plain of Sesto Fiorentino (Tuscany, Italy). Males infected by *Xenos* females are lacking in our data set, due to the scarcity of parasitized males, a secondary target of infection, and the protandrous emergence of *X. vesparum* (Hughes et al. 2004a). Wasps from each caste emerge synchronously and at specific times throughout the summer, which controls for age (Molina and O’Donnell 2008) and seasonality effects that can influence brain development. Non-parasitized and parasitized workers are easily distinguished by inspecting for extrusions between the abdominal tergites, and parasites can be identified as female or male because of the shape of their pupal sac (Fig. 1). Finally, after verifying which individuals were parasitized, their abdomens were preserved and dissected in 70% ethanol to confirm presence and sex of the parasites, and ovary development predicted for each category. We preserved each head capsule individually in a glyoxal fixative for subsequent histological sectioning (Prefer, Anatech Ltd, Battle Creek, USA).

**Figure 1.**
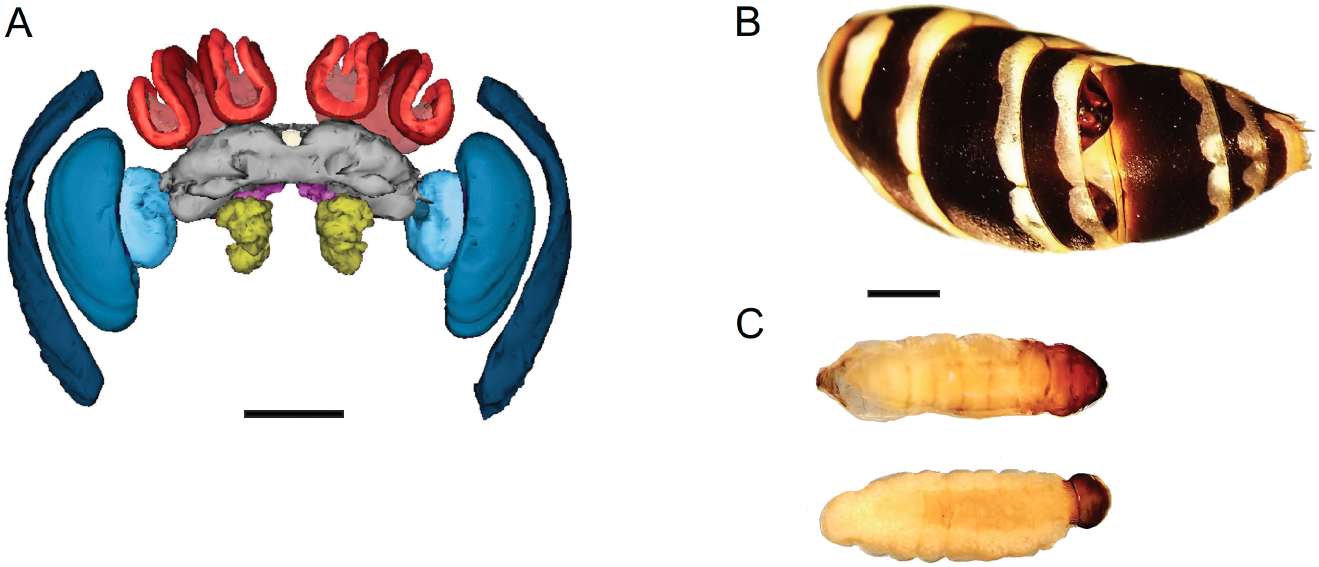
Study system: a female brain, the abdomen of a parasitized female, and *X. vesparum* parasites. A) Frontal view of a 3-D reconstructed brain of a female *Polistes dominula*. The sensory brain regions are color coded: substructures of the optic lobes (blue), antennal lobes (yellow), lip (bright red), collar (dark red) and the central complex (white). All sensory regions are normalized by the central brain (gray). For reference, the subesophageal zone is shown in bright pink and the mushroom body peduncles in light red. Scale bar = 500 μm. B) A host abdomen shows the cephalothorax of a neotenic male *X. vesparum* (top) and one female *X. vesparum* pupa (bottom) extruding from the tergites. C) Same larvae of *X. vesparum* after being dissected from the host’s abdomen: male parasite on top and female on the bottom. Figures B and C are scaled (scale bar = 2 mm).

### Histology and measurement of brain regions

We first dehydrated each head capsule with a series of increasing ethanol and acetone concentrations. We then used the established concentrations for the Embed 812 resin kit (Electron Microscopy Sciences, Hatfield, USA) to embed the head capsule while maintaining their brain dimensions, following the histology protocol for *Polistes* wasps (O’Donnell et al. 2015; Rozanski et al. 2021). The samples were moved repeatedly between an open-air rocking shaker (Thermo Fisher Scientific, Waltham, USA) and a vacuum to improve infiltration of the solvent.

Next, we placed each embedded head capsule in an individual plastic mold filled with the same concentration of resin in an oven at 60°C. After 72 hours, the resin was polymerized. We sectioned each brain in consecutive coronal sections with a thickness of 17 μm and stained the tissue with toluidine blue, to visualize clearly defined boundaries for each brain region. We photographed the consecutive brain sections for each specimen using a Canon EOS 5D Mark III mounted on a Leica DM IL LED microscope at 4x magnification, including a scale of 1000 μm.

Using the AxioVision SE64 (Zeiss, NY, USA), we outlined the area for each individual brain region. We traced the antennal lobes and the three substructures of the optic lobes: medulla, lobula and lamina. We also traced the two calyx substructures process olfactory and visual stimuli: lip and collar, respectively and the central complex. The remaining structures were grouped as the central brain, following established methods for this species (Rozanski et al. 2021). Outlining of brain regions was done blind to the category for each sample. We quantified each brain region for every other section per brain, as this method shows high accuracy (i.e., < 3.5% error for 34 μm thick sections) (Ehmer and Hoy 2000). We then determined the volume for each region by multiplying the area by the distance between sections (34 μm). We generated the 3-D brain reconstruction by using the software RECONSTRUCT (Fiala 2005). To control for the effect of head size, we measured head width. Finally, we determined the cross-sectional area of the corpora allata by measuring the diameter of one of the two glands, following the method previously used for this species (Strambi and Strambi 1973).

### Statistical analysis

We explored if differential volume in specific brain regions among phenotypes was the result of changes in allometric scaling (Eberhard and Wcislo 2011; O’Donnell et al. 2013; Ott and Rogers 2010; Seid et al. 2011; Sheehan et al. 2019; Stöckl et al. 2016). In *P. dominula*, the optic lobe represents on average 42% of the brain and may have an effect on relative neuropil scaling (Rozanski et al. 2021). Therefore, we normalized each sensory brain region by the central brain, instead of by the whole brain (Ott and Rogers 2010; Sheehan et al. 2019; Stöckl et al. 2016).

We used the allometric equation y =a*x^*β*^ for the scaling relationship between brain regions x and y. We then logarithmically transformed the estimates *β* (slope) and *α* (intercept of a regression) by using the linear equation log(y) =*β*log(x) + log(a), where log(a) = *α* (Dubois 1897; Huxley and Teissier 1936). Standardized Major (SMA) regression analyses were calculated by using the SMATR v.3 package for R (Warton et al. 2012; Warton et al. 2006).

First, we tested for a common slope among non-parasitized phenotypes as a baseline comparison, consisting of males, reproductive females and workers (H^0^ = *β_males_* = *β_reproductives_* = *β_workers_*). We implemented log-likelihood tests followed by posthoc pairwise comparisons provided in the SMATR package. Allometric scaling did not differ significantly between foundresses and gynes, we pooled them under a new category called “reproductives”. Second, we tested for a common slope among non-parasitized workers, with one female parasite and with one male parasite, and between non-parasitized and parasitized males. The volume of brain regions did not differ between male wasp parasitized by one or two male *X. vesparum*, so we also pooled them. We compared allometric changes in the whole brain with head width, central brain with whole brain, and pooled sensory regions with changes in the central brain. Finally, we explored the allometric relationship between each sensory brain region and central brain, following our established method for this species (Rozanski et al. 2021).

For categories that shared a Common Slope, we used log-likelihood tests to calculate the Slope Index (SI) for the brain region comparisons described above. The SI determined if a brain region is allometric (*β* ≠ 1), meaning that sensory brain region (y)/central brain (x) would change with size. We also used a Wald Test to calculate the Common Shift (H^0^ = equal axis among phenotypes), for any shift along the x axis. Finally, we calculated how much larger a sensory region (y) is compared to the central brain (y), by using a Grade Shift Index (GSI) to compare phenotypes (i.e., H^0^ = *α _males_* = *α _reproductives_* = *α _workers_*). The GSI reflected changes in intercept *α* (elevation) with no changes in the slope *β*. This method facilitates pairwise volumetric comparisons between phenotypes (*i.e*., e ^*α males- α* reproductives^), by implementing a Wald test. For example, if GSI > 1, males had larger volume of a brain region compared to reproductives, and if GSI < 1 the relationship would be inverse. We specify the direction of change for each of the analyzed categories in the results section and Supplementary Tables 1 and 2. Lastly, we ran a Kruskal-Wallis test with subsequent pairwise comparisons to determine corpora allata growth across castes and to test the effect of both parasite and host sex.

## RESULTS

### Investment in sensory regions by caste and sex

All brain regions, except for the central complex showed a common slope, but had unique differences in the GSI, common shift and/or SI depending on the specific region (Fig. 2., Suppl. Table 1.). Males and reproductive females had proportionally larger pooled sensory regions compared to workers (GSI = 1.056, P = 0.01 and GSI = 1.036, P = 0.006, respectively, Fig. 2). Males had proportionally smaller antennal lobes when compared to reproductive females (GSI = 0.87, P < 0.001), as an effect of both changes in elevation and a common shift (Fig. 2, Suppl. Table 1). Surprisingly, males had larger antennal lobe volume than workers (GSI = 0.926, P = 0.002). Within females, reproductives had larger antennal lobes (GSI = 1.057, P = 0.002) and calyces compared to workers (GSI = 1.042, P = 0.003). Males had larger optic lobes than reproductive females (GSI = 1.064, P = 0.001) and workers (GSI = 1.103, P = < 0.001, Suppl. Table 1). Reproductives had increased optic lobe volume compared to workers (GSI = 1.037, P = 0.02, Fig. 2). Finally, workers showed an isometric increase in the central complex (P = 0.052), in contrast to a hyperallometric reduction of this navigational brain region in reproductive females and males (Fig. 2).

**Figure 2.**
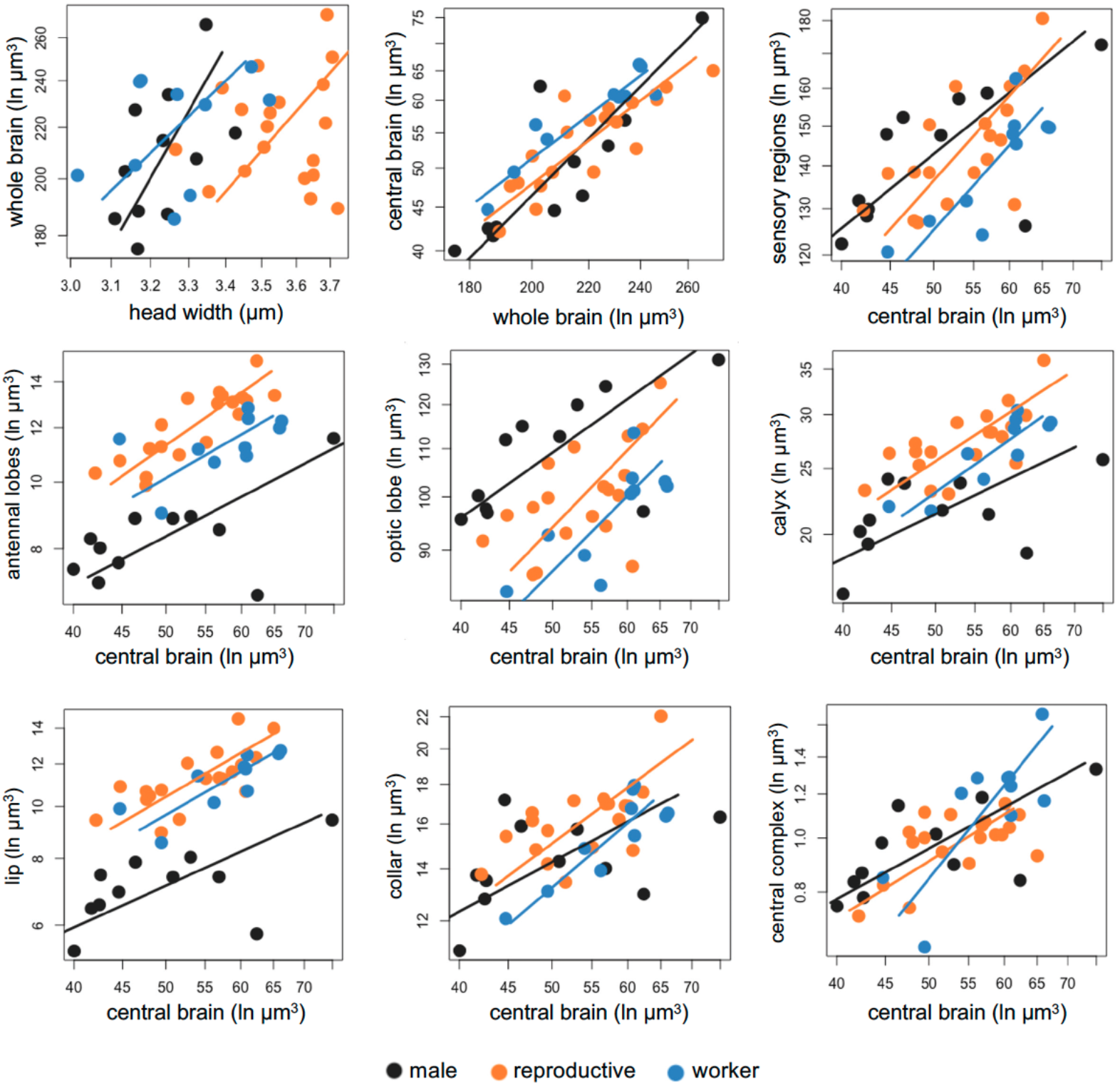
Comparison of the investment in sensory regions by sex and caste. The volume of each brain region was log-transformed, and each dot represents one individual. The males are depicted in black, reproductives (gynes and foundresses) in dark orange and workers in blue. The corresponding colored lines represent the slope for each category, and each dot represents an individual. Given that most comparisons shared a common slope, see Supplementary Table 1. for full Standardized major axis (SMA) results.

### Investment in sensory regions by parasitized and non-parasitized wasps

Surprisingly, female workers parasitized by one female or one male *X. vesparum* showed no differences in allocation of most sensory brain regions, compared to non-parasitized workers. Indeed, non-parasitized workers shared a common slope with workers with a female or a male *X. vesparum*, and no volumetric differences in the antennal lobes or the optic lobes (Fig. 3, Suppl. Table 1). However, we did find a change in the slope index of whole brain in workers parasitized by a female, compared to non-parasitized workers or parasitized by a male (P < 0.001, Fig. 3, Suppl. Table 1). Workers parasitized by one female had an isometric pattern, resulting in larger calyces (P = 0.031) and collars (P = 0.045), than non-parasitized workers and those parasitized by one male. Lastly, workers with one male parasite had a hyperallometric reduction of the central complex in comparison to non-parasitized workers and those parasitized by a female (P = 0.027, Fig. 3).

**Figure 3.**
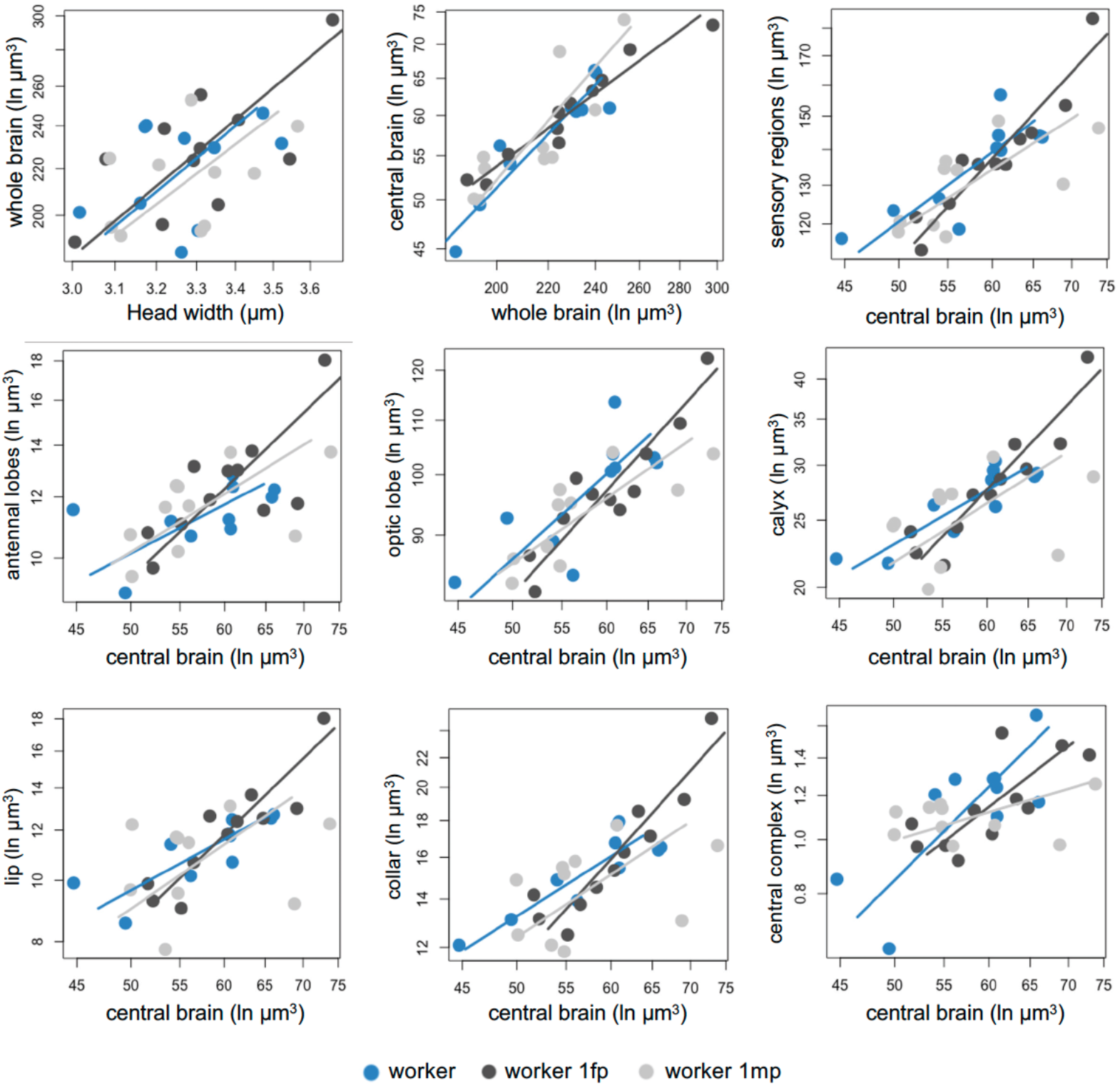
Comparison of the investment in sensory regions by parasitized and non-parasitized workers. Categories are depicted as non-parasitized workers (blue), workers parasitized by one female *X. vesparum* (dark gray) and workers parasitized by one male *X. vesparum* (light gray). The corresponding color-coded line represents the slope for each category and each dot represents an individual. Some comparisons shared a common slope, see Supplementary Table 1 for full statistical results.

In contrast, parasitized and non-parasitized males showed differential allocation towards specific brain regions. They shared a common slope and differences in grade shifts for the following brain regions: whole brain, antennal brain, lip, and central complex (Suppl. Table 2). Parasitized males had a proportionally smaller whole brains than non-parasitized males (GSI = 1.15, P < 0.001, Suppl. Table 2, Fig. 4). However, due to a common shift along the main slope axis, parasitized males had proportionally large antennal lobes (P = 0.01), lip (P = 0.001) and central complex (P < 0.001) compared to non-parasitized males (Suppl. Table 1). In contrast, the following brain regions did not show isometric growth: central brain, pooled sensory regions, optic lobes, calyces and collar (Fig. 4, Suppl. Table 2). Parasitized males showed a disproportionately reduced volume of the central brain (P = 0.02), but disproportionately large volume of pooled sensory regions (P = 0.03), optic lobes (P = 0.03), calyces (P = 0.04), and collar (P = 0.02) compared to non-parasitized males (Fig. 4, Suppl. Table 2).

**Figure 4.**
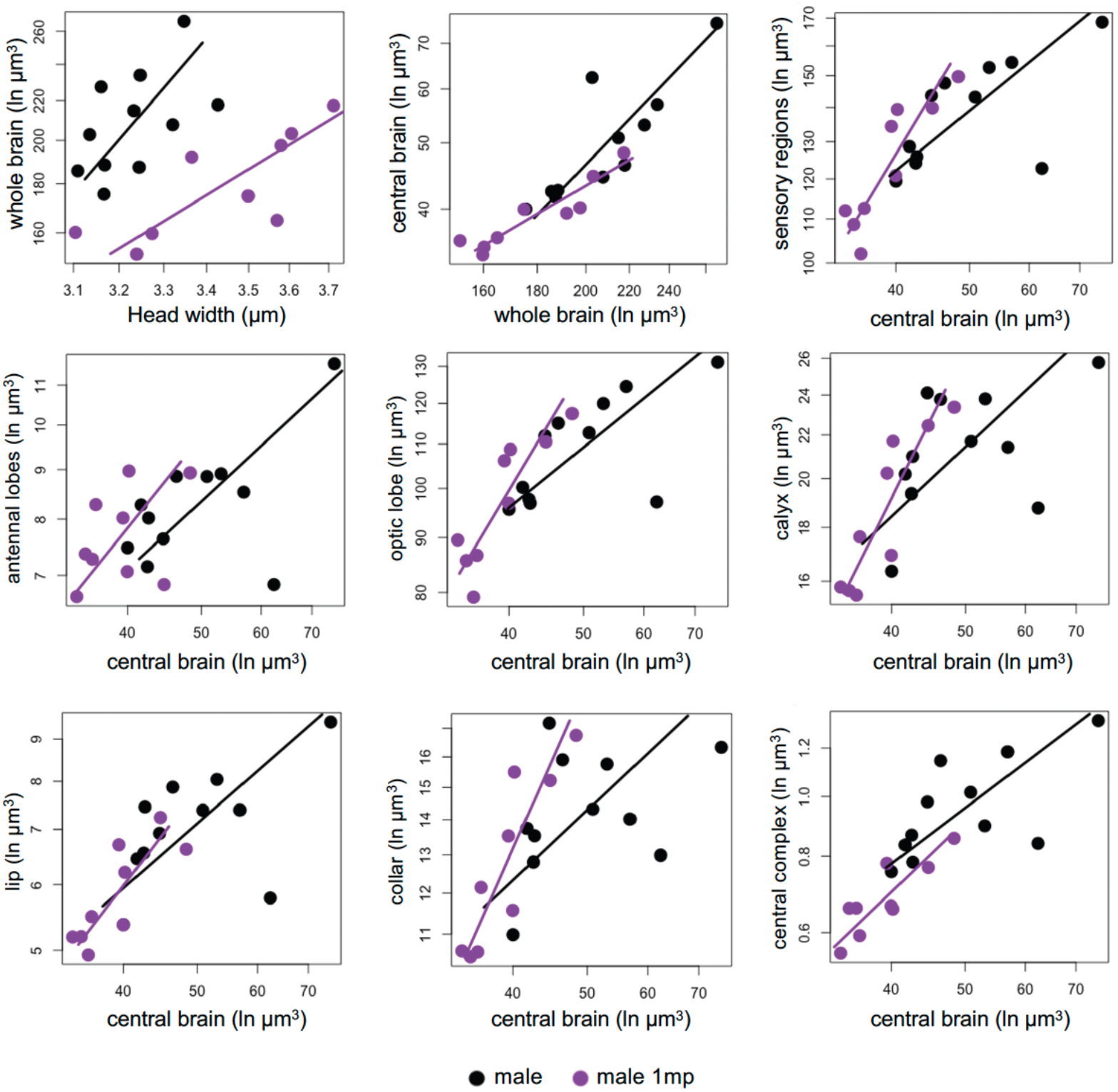
Comparison of the investment in sensory regions by parasitized (black) and non-parasitized males (gray). SMA fits are log-transformed per categories with the lines based on intercepts and slopes (black for parasitized and grey for non-parasitized males). Most volumetric comparisons did not share a common slope, see Supplementary Table 2 for full SMA tests.

### Corpora allata development according to sex, caste and parasitism

The corpora allata were significantly smaller in all males compared to females (χ^2^ = 46.86, df = 6, P < 0.001, Fig. 5). As expected, foundresses had larger corpora allata compared to gynes and workers. Posthoc pairwise tests revealed showed a trend in reduction of the corpora allata in parasitized workers compared to non-parasitized workers. However, gland size did not differ according to parasite sex (P = 0.39), or between parasitized and non-parasitized males (P = 0.98).

**Figure 5.**
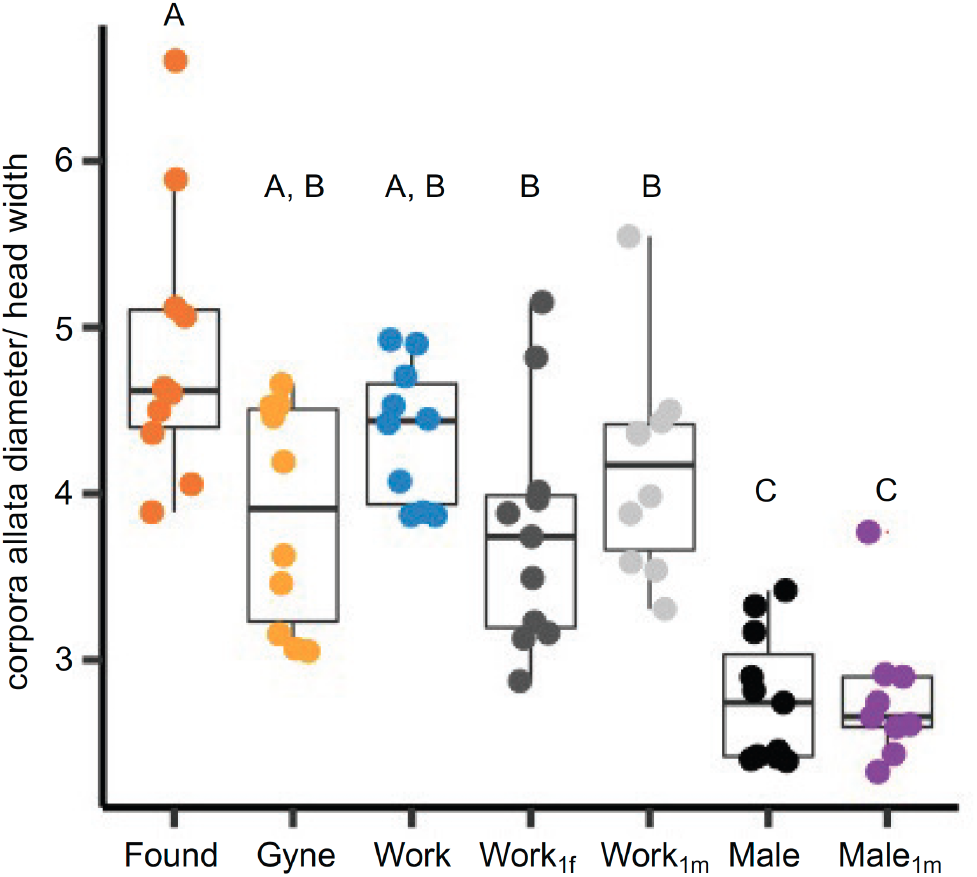
Corpora allata size in reproductive females, workers, males, parasitized workers and parasitized males. The cross-sectional area of the corpora allata was calculated by measuring its diameter in microns and normalized with head size. Wasp categories are color coded: found (foundresses), gynes, work (workers), work_1fm_ (workers with 1 female parasite), work_1m_ (worker with 1 male parasite), males and male_1m_ (male with 1 male parasite).

## DISCUSSION

We provide several lines of evidence supporting the focal hypothesis that brain plasticity facilitates differential sensory needs and life histories within the same species. First, reproductive females also had larger calyces compared to worker females, reflecting sensory needs associated to division of labor, implying a life history-based plasticity of otherwise genetically shared backgrounds in female wasps. Furthermore, genetically distinct (haploid) males and (diploid) females showed a consistent and significant tradeoff in volume of the optic and antennal lobes. Second, we provide novel evidence for the effect of the *Xenos* parasite in neural investment by female and male hosts. Surprisingly, non-parasitized and parasitized workers show moderate volumetric differences in brain sensory regions, while parasitized males showed a more drastic effect in allocation of neural tissue compared to non-parasitized workers. Overall, our results are consistent with differential investment in brain regions being advantageous across social wasp species (O’Donnell et al. 2011), while it can also be driven by parasitic manipulation, which may be potentially maladaptive for the host.

The observed neural tradeoffs reflect the distinct life cycles of *P. dominula* males and females, similarly to previous studies in social Hymenoptera (reviewed in Beani et al. 2014). Males leave their nest within a few days after emergence and gather at distinct leks to increase their mating opportunities (Beani 1996). When attempting to mate, males can visually distinguish between females and competing males, and between workers and gynes (Cappa et al. 2013; da Silva et al. 2021; de Souza et al. 2017). Therefore, higher investment in the optic lobes may facilitate detection and discrimination between potential mates or male intruders in their defended territories (Beani et al. 2014). Males have smaller antennal lobes, which is likely due to experiencing less complex olfactory stimuli, as they do not engage in frequent social interactions in the colony. In contrast, reproductive females have proportionally larger antennal lobes, lips and collars, which is consistent with other studies that show sensory needs associated to division of labor, interactions among nestmates, learning and memory (Ehmer and Hoy 2000; Gronenberg et al. 1996; Jernigan et al. 2021; Mertes et al. 2021; O’Donnell et al. 2011; Rozanski et al. 2021; Uy et al. 2021). Thus, the social environment of female wasps has a wider range of chemical and sensory processing cues compared to males (Beani et al. 2014). Within females, reproductives had proportionally larger calyces than workers, which coincides with division of labor in these social wasps (Molina and O’Donnell 2008; O’Donnell et al. 2007). Future foundresses consistently engage in social interactions both within the colony and during winter aggregations, utilizing visual and chemical cues towards recognition (Cini et al. 2019; Dani et al. 2001). In contrast, most workers spend less time interacting with foundresses and brood on the nest, and allocate more time performing tasks such as foraging for prey and building material (Gamboa et al. 1978).

We confirmed reduced corpora allata in parasitized workers, when compared to non-parasitized ones and more drastically to reproductive females (Strambi and Strambi 1973). However, contrary to our expectations, parasites have milder effects in the brain architecture of workers. Parasitized workers lose most of their social behavior and aggregate on specific plants where they perform site-fidelity and territorial defense of resources, allowing the completion of the parasite life cycle (Beani et al. 2018). This active life style most likely requires intact sensory processing and cognitive abilities. By selectively acting on the neuroendocrine function of the corpora allata (Strambi and Strambi 1973), the parasite may specifically disrupt the host social behavior while maintaining the sensory machinery needed to survive. Thus, the parasite has strong effects in castrating the female host and in inducing her social aberrant behavior, but moderate effects on allocation of brain regions.

Noticeably, workers parasitized by one *X. vesparum* female showed larger calyces than non-parasitized workers and those parasitized by one male. Interestingly, workers parasitized by one female overwinter and resemble the behavioral and physiological phenotype of overwintering gynes, while workers parasitized by a male die at the end of summer like non-parasitized ones (Beani et al. 2021). Male parasites also induced a hyperallometric reduction of the central complex, compared to non-parasitized workers or workers parasitized by one female. This difference in the central complex, which is mainly implicated in spatial navigation, is not consistent with the lack of differences in behavior between workers infected by the two sexes.

In contrast, male parasites had a more drastic effect in the brain architecture of *P. dominula* males, which act as secondary hosts. Parasitized males had significantly smaller whole brains and central brains than non-parasitized males. They also showed a significant increase in the volume of several sensory brain regions, including the antennal and optic lobes, and two substructures of the calyx: lip and collar. Remarkably, neuroendocrine manipulation does not seem to occur in parasitized males, as they can still develop their testes and attempt to mate with females (Beani et al. 2017; Cappa et al. 2014). The inability to hinder male reproduction may likely result in a tradeoff with brain manipulation, which is an expensive tissue to produce (Keesey et al. 2020; Niven and Laughlin 2008).

Overall, our results demonstrate that brain plasticity is associated to sensory needs in males and within female castes of *P. dominula*, but that parasitic manipulation can also drive differential investment of brain regions depending on both host and parasite sex. Intriguingly, parasitized workers show a strong manipulation effect of the parasite on caste determination, lipid storage, and prolonged lifespan in parasitized females (Beani et al. 2021), but more dampened effects on allocation of brain tissue. In turn, the reproductive apparatus and behavior of parasitized males are essentially unaffected, but they experience stronger volumetric changes in brain regions. However, the parasite may be relying on other manipulation mechanisms, without driving evident neuroanatomical changes in its female hosts (Libersat et al. 2018). For instance, *X. vesparum* can drive changes in gene expression of a worker towards a gyne-like pattern; thus, the parasite is manipulating the transcriptomic plasticity of the caste system (Geffre et al. 2017). Future research in organisms that show natural plasticity in behavioral and sensory needs will provide further opportunities to explore the neural mechanisms underlying parasitic manipulation.

## Supporting information

Gandia_etal_Suppl_Material

## ACKNOWLEDGEMENTS

We thank Jamie Brunworth for assistance in quantifying brain regions. William Searcy, Al Uy and members of the Uy lab provided useful feedback in early versions of the manuscript. KMG, MEH and FMKU were supported by a National Academies Keck Future Initiatives Grant (NAKFI). FC, DB and LB were supported by University of Florence funds. MEH is a visiting fellow at the Wissenschaftskolleg zu Berlin. Samples were imported from Italy into the USA via USDA permit #128388.

## AUTHOR CONTRIBUTIONS

LB and FC collected the wasp and parasite field samples. KMG performed histological preparation of specimens and collected volumetric data. FMKU analyzed the data. KMG, LB, FC and FMKU contributed to the first draft of the manuscript. All authors provided input to concept and design of this project, data interpretation, along with reviewing and editing the final manuscript. LB and FMKU shared senior authorship.

