## Supplementary material for "Caste, sex, and parasitism influence brain plasticity in a social wasp": Gandia_etal_Suppl_Material

**SUPPLEMENTARY TABLE 1**

Standardized Major Axis regressions (SMA) exploring the allometric changes in brain plasticity according to caste, sex, and parasitism in the social wasp *Polistes dominula*. We compared the role of allometry in the volume of whole brain (WB), central brain (CB), combined sensory brain regions (SENS), the antennal lobes (ALs), optic lobes (OLs), calyx (CAL) and its substructures lip (LI), collar (CO), and the central complex (CX). We used the Grade Shift Index (GSI = $e^{\alpha\ host\ -\ \alpha\ par}$) to determine changes in elevation (intersect) between two experimental categories by comparing the scaling relationship between each log-transformed sensory region and the CB. The categories included males (male), reproductive females (rep), non-parasitized workers (work), workers parasitized by 1 female (wpf) or 1 male (wpm) *X. vesparum*. If GSI > 1, a specific brain region is larger in the first category of the pairwise comparisons (*i.e.*, GSI > 1 reflects male > rep, and GSI < 1 reflects male < rep). Significance for the GSI is reported as: P > 0.05, P < 0.05, P < 0.01. The Slope Index (SI) determines if the allometric scaling of each sensory brain region to the central brain deviates from $\beta$ = 1. We implemented SMA tests recommended by the SMATR 3 R package (Warton et al. 2012; Warton et al. 2006) .

**SEX & CASTE**

| | Common Slope | | Common Elevation | | | | | Common Shift | | | | Isometry | | |
|---|---|---|---|---|---|---|---|---|---|---|---|---|---|---|
| | | | | | GSI | | | | Posthoc comparisons | | | | | |
| **Brain region** | **Log likelihood** | ***P*** | **Wald test** | ***P*** | **male-rep** | **male-work** | **rep-work** | **Wald test** | **male-rep** | **male-work** | **rep-work** | **Log likelihood** | ***P*** | **SI** |
| WB | 2.07 | 0.36 | 15.18 | <0.001 | 1.10 | 0.99 | 0.90 | 17.70 | <0.001 | 0.72 | 0.03 | 0.54 | <0.001 | 1.38 |
| CB | 1.69 | 0.44 | 10.17 | 0.01 | 1.00 | 0.97 | 0.97 | 2.49 | 0.56 | 0.31 | 0.84 | 0.57 | <0.001 | 1.44 |
| SENS | 1.18 | 0.55 | 13.02 | 0.00 | 1.02 | 1.06 | 1.04 | 1.03 | 0.78 | 0.72 | 0.98 | -0.47 | 0.00 | 1.01 |
| ALs | 1.06 | 0.59 | 44.93 | <0.001 | 0.87 | 0.92 | 1.06 | 20.42 | <0.001 | <0.001 | 1.00 | 0.33 | 0.04 | 1.32 |
| OLs | 1.38 | 0.50 | 25.88 | <0.001 | 1.06 | 1.10 | 1.04 | 0.15 | 0.98 | 1.00 | 0.98 | 0.16 | 0.01 | 0.73 |
| CAL | 1.22 | 0.45 | 16.57 | <0.001 | 0.93 | 0.96 | 1.04 | 12.16 | 0.00 | 0.01 | 0.97 | 0.06 | 0.72 | 1.04 |
| CO | 1.63 | 0.44 | 12.20 | 0.00 | 0.98 | 1.03 | 1.05 | 4.97 | 0.11 | 0.19 | 0.99 | 0.20 | 0.22 | 0.85 |
| LI | 0.58 | 0.75 | 42.08 | <0.001 | 0.85 | 0.87 | 1.03 | 27.07 | <0.001 | <0.001 | 0.88 | 0.52 | <0.001 | 1.61 |
| CX | 5.89 | 0.05 | | | | | | | | | | | | |

**PARASITISM IN WORKERS**

| | Common Slope | | Common Elevation | | | | | Common Shift | | | | Isometry | | |
|---|---|---|---|---|---|---|---|---|---|---|---|---|---|---|
| | | | | | GSI | | | | Posthoc comparisons | | | | | |
| **Brain region** | **Log likelihood** | ***P*** | **Wald test** | ***P*** | **w-wpf** | **w-wpm** | **wpf-wpf** | **Wald test** | **work-wpf** | **work-wpm** | **wpf-wpm** | **Log likelihood** | ***P*** | **SI** |
| WB | 0.06 | 0.97 | 0.77 | 0.68 | 0.99 | 1.01 | 1.02 | 0.82 | 0.87 | 0.99 | 0.75 | 0.72 | <0.001 | 2.21 |
| CB | 6.36 | 0.04 | | | | | | | | | | | | |
| SENS | 4.36 | 0.11 | 0.74 | 0.69 | 1.01 | 1.01 | 1.00 | 1.22 | 0.87 | 0.95 | 0.62 | 0.20 | 0.29 | 0.88 |
| ALs | 2.69 | 0.26 | 0.46 | 0.79 | 0.98 | 0.99 | 1.00 | 1.72 | 0.53 | 1.00 | 0.65 | 0.13 | 0.47 | 1.11 |
| OLs | 2.78 | 0.25 | 1.34 | 0.51 | 1.01 | 1.01 | 1.00 | 1.03 | 0.93 | 0.94 | 0.67 | 0.33 | 0.03 | 0.81 |
| CAL | 6.91 | 0.03 | | | | | | | | | | | | |
| CO | 6.16 | 0.05 | | | | | | | | | | | | |
| LI | 4.42 | 0.11 | 0.11 | 0.95 | 1.00 | 1.01 | 1.01 | 1.51 | 0.76 | 0.98 | 0.55 | 0.40 | 0.02 | 1.40 |
| CX | 7.16 | 0.03 | | | | | | | | | | | | |

**SUPPLEMENTARY TABLE 2**

Standardized Major Axis regressions (SMA) testing for the allometric changes in the volume of sensory brain regions in non-parasitized and parasitized males. Brain regions are depicted in the same manner as Suppl Table 1: (WB), central brain (CB), combined sensory brain regions (SENS), the antennal lobes (ALs), optic lobes (OLs), calyx (CAL) and its substructures lip (LI), collar (CO), and the central complex (CX). Most brain regions did not share a common slope, and instead showed disproportionate increase or reduction compared to the CB. For regions that shared a common slope, if GSI > 1, the region is proportionately larger in non-parasitized males. If GSI <1, the region is larger in parasitized males.

| | Common Slope | | Common Elevation | | | Common Shift | | Isometry | | |
|---|---|---|---|---|---|---|---|---|---|---|
| **Brain region** | **Log likelihood** | ***P*** | **Wald test** | ***P*** | **GSI** | **Wald test** | **P** | **Log likelihood** | ***P*** | **SI** |
| WB | 2.71 | 0.09 | 29.43 | <0.001 | 1.14 | 0.01 | 0.89 | 0.75 | <0.001 | 2.64 |
| CB | 5.17 | 0.02 | | | | | | | | |
| SENS | 4.49 | 0.03 | | | | | | | | |
| ALs | 0.40 | 0.52 | 2.46 | 0.11 | 0.95 | 5.78 | 0.010 | 0.46 | 0.03 | 0.65 |
| OLs | 4.67 | 0.03 | | | | | | | | |
| CAL | 4.31 | 0.03 | | | | | | | | |
| CO | 15.26 | <0.001 | | | | | | | | |
| LI | 1.15 | 0.28 | 0.49 | 0.48 | 0.97 | 9.80 | 0.001 | 0.20 | 0.39 | 0.87 |
| CX | 0.52 | 0.47 | 1.12 | 0.28 | 1.03 | 15.26 | <0.001 | 0.25 | 0.27 | 1.16 |
